# An embryonic zebrafish model to screen disruption of gut-vascular-barriers upon exposure to ambient ultrafine particles

**DOI:** 10.1101/2020.10.28.359976

**Authors:** Kyung In Baek, Yi Qian, Chih-Chiang Chang, Ryan O’Donnell, Ehsan Soleimanian, Constantinos Sioutas, Rongsong Li, Tzung K. Hsiai

**Affiliations:** Departments of Bioengineering and Medicine, University of California, Los Angeles, California, 90095; Department of Cardiology, The Second Affiliated Hospital, School of Medicine, Zhejiang University, China; Department of Civil and Environmental Engineering, University of Southern California, Los Angeles, California, 90089; College of Health Sciences and Environmental Engineering, Shenzhen Technology University, Shenzhen, China; Division of Cardiology, Department of Medicine, School of Medicine, University of California, Los Angeles, California, 90095; Veterans Affairs Greater Los Angeles Healthcare System, Department of Medicine, Los Angeles, California, 90073

**Keywords:** Ultrafine particles, zebrafish, micro-gavage, Notch signaling, gut vascular barrier

## Abstract

Epidemiological studies have linked exposure to ambient particulate matter (PM) with gastrointestinal (GI) diseases. Ambient ultrafine particle (UFP) are the redox-active sub-fraction of PM2.5, harboring elemental and polycyclic aromatic hydrocarbons from urban environmental sources including diesel and gasoline exhausts. The gut vascular barrier (GVB) regulates paracellular trafficking and systemic disseminations of ingested microbes and toxins. Here, we posit that acute UFP ingestion disrupts the integrity of the intestinal barrier by modulating intestinal Notch activation. Using zebrafish embryos, we performed micro-gavage with the FITC-conjugated dextran (FD10, 10 kDa) to assess the disruption of GVB integrity upon UFP exposure. Following micro-gavage, FD10 retained in the embryonic GI system, migrated through the cloaca. Conversely, co-gavaging UFP increased transmigration of FD10 across the intestinal barrier, and FD10 fluorescence occurred in the venous capillary plexus. Ingestion of UFP further impaired the mid-intestine morphology. We performed micro-angiogram of FD10 to corroborate acute UFP-mediated disruption of GVB. Transient genetic and pharmacologic manipulations of global Notch activity suggested Notch regulation of the GVB. Overall, our integration of a genetically tractable embryonic zebrafish and micro-gavage technique provided epigenetic insights underlying ambient UFP ingestion disrupts the GVB.

**Graphic Abstract:** 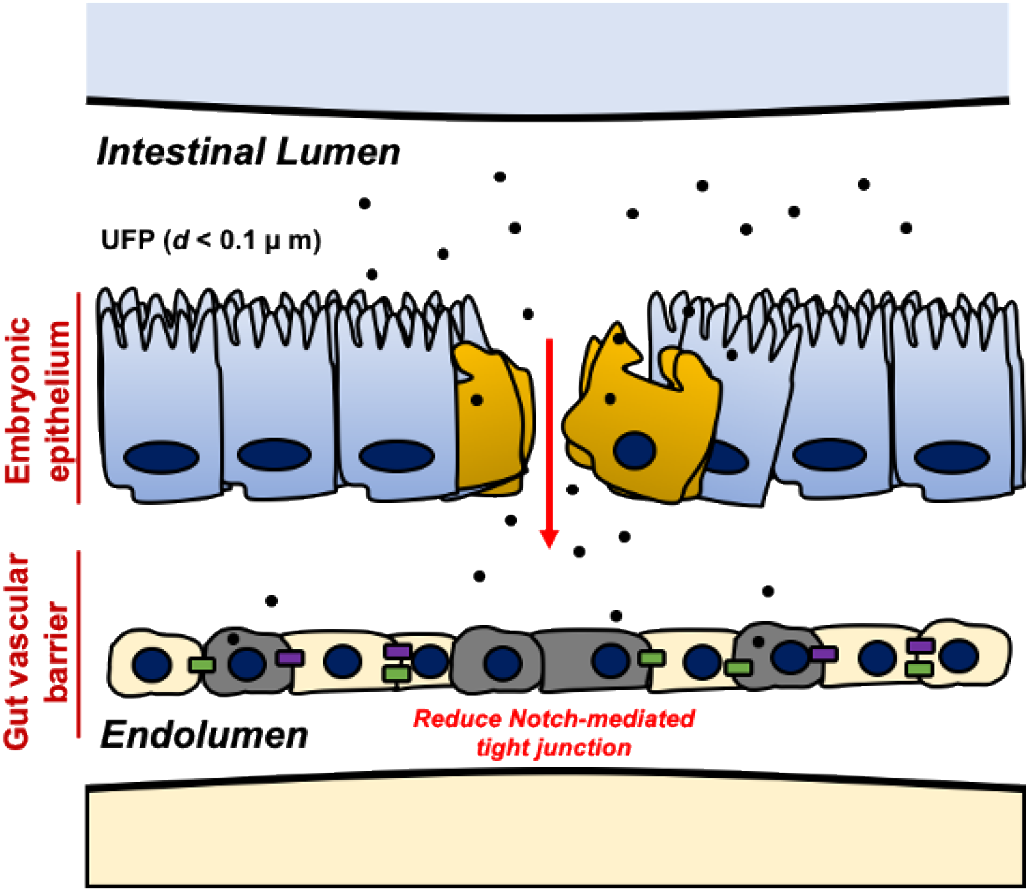

## 1. Introduction

Ultrafine particles (UFP, *d_p_* < 0.1 to 0.2 μm in diameter), comprised of a mixture of heavy transition metals and redox cycling organic chemicals, are redox-active components of ambient particulate matter (PM, *d_p_* < 2.5 μm).[1,2] Recent epidemiological studies have supported the link between UFP exposure and gastrointestinal (GI) diseases such as inflammatory bowel disease.[3] While the cardiopulmonary system remains the main entry point for ambient UFP exposure, UFP are orally ingested via contaminated food and water supplies for intestinal pro-inflammatory potentials. [4–7] Dietary UFP, including titanium dioxide nanoparticles used as food additives and aluminosilicate minerals in drinking water, are absorbed by intestinal epithelial lymphocytes, potentiating pro-inflammatory cytokines and T-cell proliferation.[8,9] Inhaled UFP increases colonic inflammation and facilitates intestinal release of fatty acids and pro-inflammatory mediators as a result of bronchial mucocilliary removal to the oropharynx.[10] While intestinal inflammatory responses suppress mucosal stability and subsequent integrity of the intestinal epithelial barrier, the epigenetic cues underlying ambient UFP exposure and the barrier integrity remain elusive.[11]

The gut vascular barrier (GVB) constitutes both intestinal epithelial and vascular endothelial barriers that regulate dissemination of microbes and toxins from the intestinal tract to systemic circulation.[12–14] Analogous to the blood-brain barrier, a layer of endothelial cells along with surrounding glial cells and pericytes, the GVB develops cellular junctions to form a functional barrier, regulating paracellular trafficking (< 4 kDa) into the vascular endolumen.[13,14] Junctional complexes in an endothelial layer including tight junctions (TJ), occludin, zonula occludens-1 (ZO-1), claudin, and adherens proteins, including vascular endothelial-cadherin and β-catenin characterize the integrity of the GVB. Furthermore, complementary mechanisms of endothelial transcytosis and GVB homeostasis share similarities with the blood-brain barrier.[15] Microbial pathogens such as Salmonella typhimurium or celiac dysbiosis modulate the Wnt3a/β-catenin signaling pathway to reduce enteric TJ expression, and dismantle the GVB for diffusion of intestinal pro-inflammatory mediators.[14,16] While acute UFP exposure is reported to increase both colonic epithelial and endothelial permeability *in vitro*, in conjunction with an altered diversity of gut microbiota in low density lipoprotein receptor-null mice (ldlr^-/-^), the molecular cues whereby UFP ingestion disrupts the GVB remain elusive.[10,17]

In this context, we posit that acute UFP ingestion via micro-gavage disrupt the GVB by inhibiting Notch-dependent TJ expression. To visualize disruption of GVB, we sought to use the embryonic zebrafish (*Danio rerio*) model for its transparent anatomic features and conserved genetic systems during organogenesis.[18] We demonstrated a micro-gavage technique in order to deliver UFP directly to the embryonic intestinal bulbs for rapid screening the disruption of the GVB. In FITC-conjugated dextran (FD10, 10 kDa)-gavaged controls, FD10 remained in the intestinal bulbs and the mid-intestine migrating only though the cloaca. At 7 hours post gavage (hpg), co-gavaging UFP (25-50 μg·mL^-1^) attenuated the intestinal barrier and global development of the GI tract; thereby, allowing for the transmigration of FD10 from the intestinal lumen into the anterior and caudal capillary venous plexuses (CVP). Micro-angiogram of FD10 via the common cardinal vein (CCV) conferred that UFP exposure disrupted the GVB. In addition, co-gavaging a disintegrin and metalloproteinase domaincontaining protein 10 (Adam10) inhibitor to inhibit extracellular proteolysis of Notch receptors mimicked UFP gavage, whereas transient overexpression of Notch signaling via Notch intracellular cytoplasmic domain (*NICD*) mRNA injection restored UFP-mediated GVB. As a corollary, UFP down-regulated transcription of cytoplasmic zonula occludens1 (Zo1), and the transmembrane claudin 1 (Cldn1), and the Notch target, Hairy and enhancer of split-1 (Hes1) in cultured endothelial cells. Overall, the integration of a genetically tractable embryonic zebrafish model with micro-gavage and optical imaging techniques provides the high-throughput screening insights into epigenetic and genetic interaction underlying UFP-mediated disruption of the GVB.

## 2. Materials and Methods

### Zebrafish maintenance and study approval

All zebrafish experiments were performed in compliance with UCLA Institutional Animal Care and Use Committee (IACUC) protocol (A3196-01).

### Collection, extraction, and chemical analysis of ultrafine particles (UFP)

Ambient ultrafine particulate matter (UFP particles with aerodynamic diameter < 0.2 μm) was collected on PTFE membrane filters (20 × 25 cm, 3.0 μm pore size, PALL Life Sciences, USA) at the University of Southern California’s Particle Instrumentation Unit using a high-volume sampler (with a flow rate of 250 liters per minute, lpm) connected to a PM2.5 pre-impactor for separation and collection of PM2.5. The loaded filter samples were then extracted in Milli-Q water (Millipore A-10, EMD Millipore, Billerica, MA, USA) following 1 hour of sonication, in which the amount of extracted PM2.5 via sonication was determined by subtracting the pre-extraction and post-extraction weights of the filters using a high precision (± 0.001 mg) microbalance (MT5, Mettler Toledo Inc., Columbus, OH). Following filter extraction, the highly concentrated aqueous suspensions of PM2.5 were diluted to provide the required slurry concentration and stored at −20 °C for experiments. Precise descriptions of sample collection and extraction can be found in Taghvaee *et al*.[19] The aqueous suspensions of PM2.5 were chemically analyzed for total organic carbon (TOC), water-soluble inorganic ions, and metal elements by Wisconsin State Laboratory of Hygiene. In summary, TOC content of the samples was quantified by means of a Sievers 900 TOC analyzer. [20,21] In addition, ion chromatography was employed to determine water-soluble inorganic ions,[22] whereas the metal and trace elements were quantified by inductively-coupled plasma mass spectroscopy analysis. [23] Chemical compositions of UFP are characterized in our previous publication.[24]

### Zebrafish culture, micro-injection, and micro-gavage assay

Zebrafish embryos were harvested from natural mating consisting of a mixture of males and females at the UCLA Zebrafish Core Facility. Embryos were cultivated at 28.5 °C in fresh standard E3 medium supplemented with 0.05% methylene blue (Sigma Aldrich, MO) and 0.003% phenylthiourea (PTU, Sigma Aldrich, MO) to suppress fungal outbreak and to inhibit melanogenesis. Ectopic overexpression of global Notch signaling pathway was performed by micro-injecting *NICD* mRNA (10-20 pg·nL^-1^) as previously described.[25] At 2 days post fertilization (dpf), embryos were manually dechorionated for micro-gavage assay.[26] To perform micro-gavage, transgenic *Tg*(*flk1: mCherry*) embryos were immobilized with neutralized tricaine (Sigma Aldrich, MO) and oriented in 1% low melting agarose (Thermofisher, MA). A solution of FD10 (Sigma Aldrich, MO) containing 0.05% phenol-red dye (Sigma Aldrich, MO) was micro-gavaged into the anterior intestinal bulb without damaging the esophagus, the swim bladder, or the yolk sac (**Figure 1D**). UFP (25-50 μg·mL^-1^), Adam10 inhibitor (5 μM, GI254023X, Sigma Aldrich, MO), or EDTA (20 mM, Sigma Aldrich, MO) were homogenized in the FD10 solution respectively for micro-gavage. At 7 hpg, distribution of FD10 in the AVP and CVP were imaged with dual channel confocal (Leica SP8, Germany) or inverted fluorescence microscopy (Olympus, IX70) previously described. [24,27]

**Figure 1.**
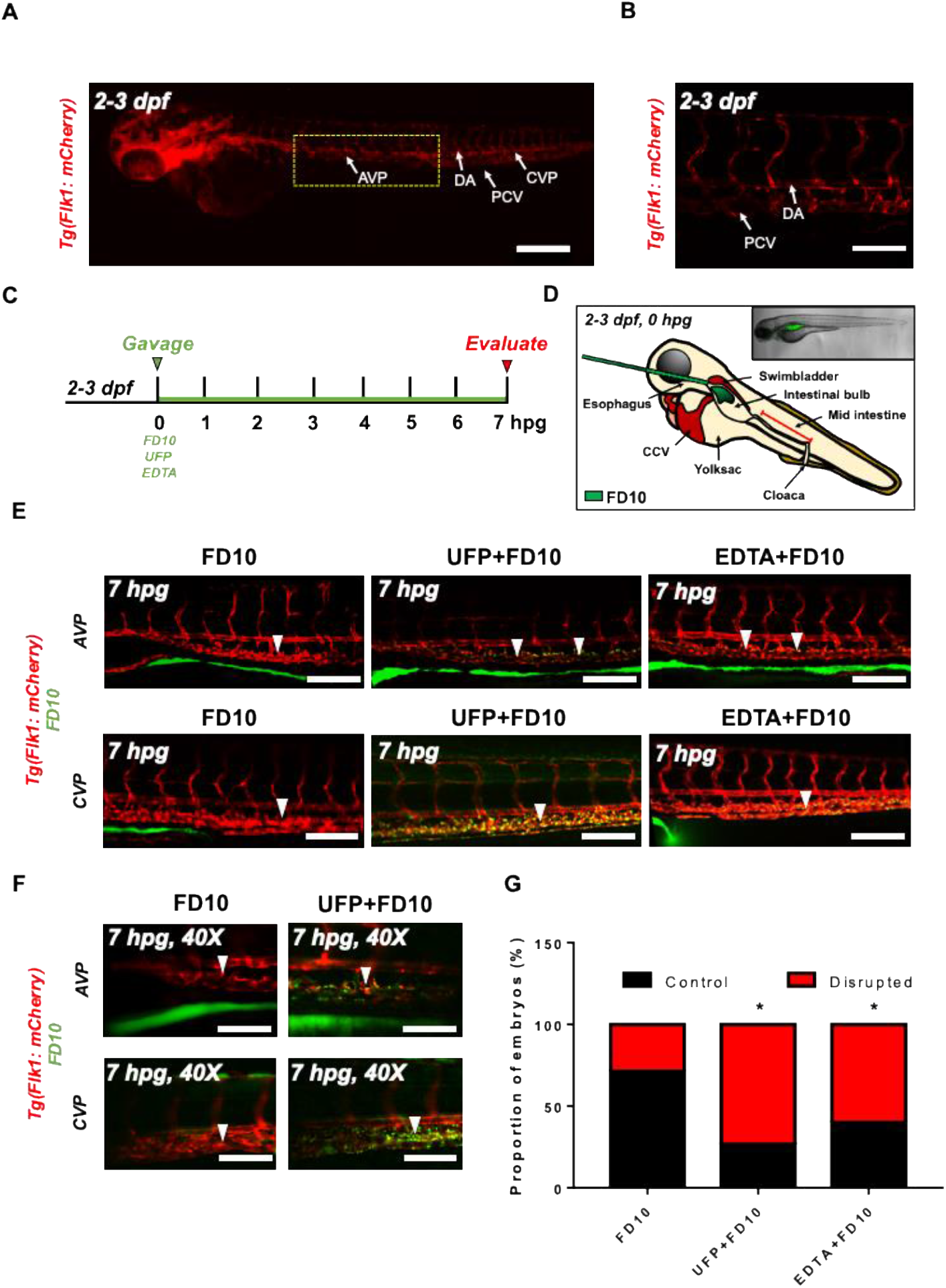
Acute UFP ingestion disrupts intestinal epithelial barrier integrity. Transgenic *Tg*(*flk1: mCherry*) zebrafish embryos at 2-3 dpf were micro-gavaged with FITC-conjugated dextran (FD10, 10 kDa). **(A-B)** Anatomy of endothelial vasculature in the *Tg*(*flk1: mCherry*) embryo. DA: Dorsal aorta; PCV: Posterior caudal vein; AVP: Anterior venous capillary plexus; CVP: Caudal vein capillary plexus; Scale bar: 32 μm. **(C)** Experimental design: At 2 dpf, embryos were randomly chosen for micro-gavage with FD10 solution with or without UFP or EDTA at 20 mM. Intestinal barrier integrity and translocation of FD10 to vascular endolumen (*flk1*^+^) were evaluated at 7 hours post gavage (hpg). **(D)** A schematic representation of micro-gavage in an embryonic GI tract. FD10 solution was micro-gavaged in the intestinal bulb without disrupting the esophagus, swim bladder and yolk sac. (E) Representative images of the AVP and CVP at 7 hpg. In FD10 gavaged-controls, FD10 remained only in the intestinal bulb and mid-intestine. In contrast, co-gavaging FD10 with UFP or EDTA accumulated FD10 in the AVP and CVP (white arrow heads). Scale bar: 20 μm. **(F)** Percentage of embryos exhibiting endoluminal FD10 fluorescence (* *p < 0.05* vs. FD10, n=10 per group).

### Micro-angiogram via common cardinal vein (CCV)

Immobilized *Tg*(*flk1: mCherry*) embryos were placed on a 3% agar plate to perform the micro-angiography. A mixture of the gavage solution (FD10 solution and 0.05% phenol-red dye) was injected into the zebrafish CCV as previously described. [28,29] Micro-angiogram was verified by imaging FD10 fluorescence in the DA and PCV. At 1 hour post injection, injected embryos were embedded in 1% low melting agarose for fluorescence imaging. Distribution of the FD10 in the AVP and CVP was evaluated under inverted fluorescence microscope (Olympus, IX70).

### Human aortic endothelial cell (HAEC) culture and quantitative real-time polymerase chain reaction (qRT-PCR) analyses

For qRT-PCR analyses, HAEC (Cell Applications, CA) between passages 4 and 10 were cultivated on 0.1% bovine gelatin-coated plates (Midsci, MO) at 37 °C and 5% CO2. EC growth medium (Cell Applications, CA) was supplemented with 5% fetal bovine serum (FBS, Life technologies, NY) and 1% penicillin-streptomycin (P/S, Life Technologies, NY) to promote cultivation. Confluent HAEC monolayers were exposed to UFP, Adam10 inhibitor, Iwr1 (10 μM, Sigma Aldrich, MO), or lithium chloride (LiCl, 20 mM) respectively in neutralized M199 media (Life Technologies, NY). Total RNA was collected following 6 hours of treatment. RNA purification and reverse transcription were performed as previously described.[27] qPCR master-mix (Applied Biological Materials Inc., Canada) was used for PCR amplification. mRNA expression was normalized to human actin expression. Sequences of primers are listed in **Table 1**.

**Table 1.**
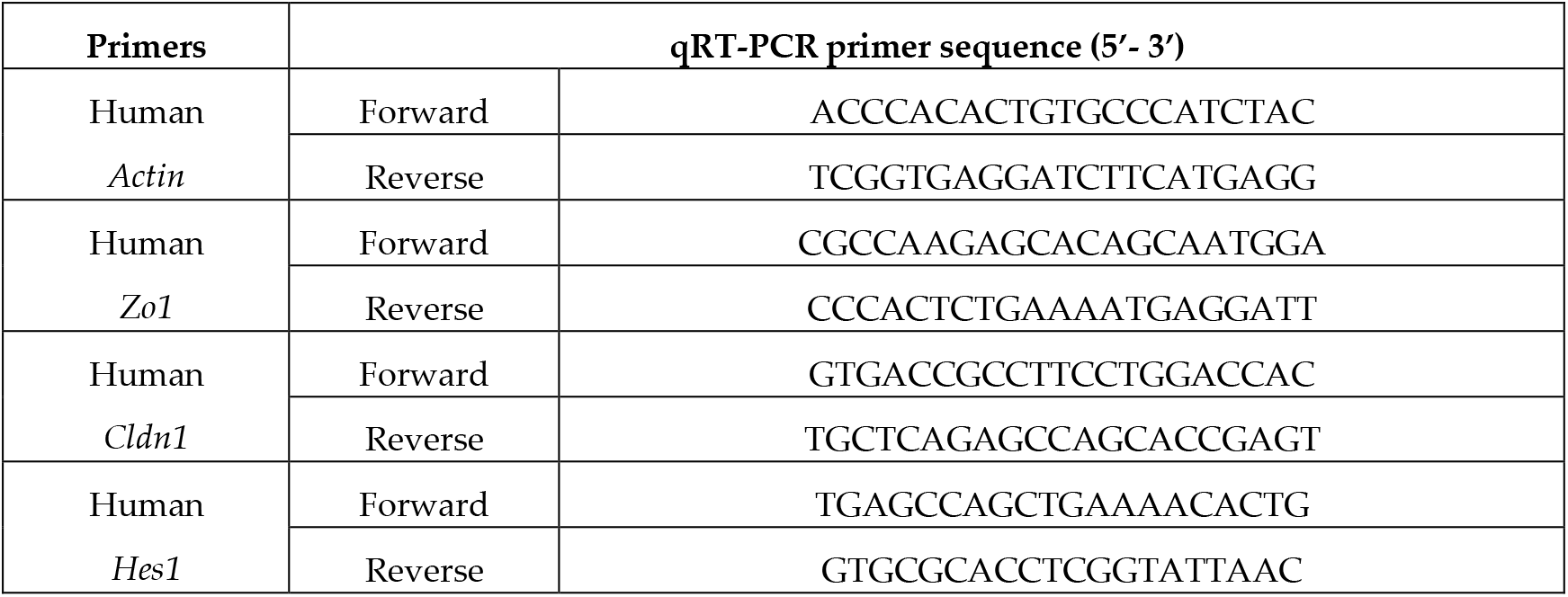
Sequencing Information of qRT-PCR primers.

### Statistics

Data were expressed as mean ± standard deviation and compared among separate experiments. Unpaired two-tail t test and 2-proportion z-test were performed for statistical comparisons between 2 experimental conditions. P values < 0.05 were considered significant.

## 3. Results

### 3.1. Acute UFP ingestion disrupts intestinal barrier integrity and maturation of GI tract

To assess whether acute UFP ingestion disrupts the embryonic intestinal barrier, FD10 suspension containing UFP (25 μg·mL^-1^) or the positive control EDTA (20 mM) was micro-gavaged to transgenic *Tg*(*flk1: mCherry*) zebrafish embryos between 2 and 3 days post fertilization (dpf) (**Figure 1A-C**). Modulated intestinal permeability and the barrier integrity as denoted by co-localization of FD10 fluorescence and vascular endothelium (*flk1^+^*) was evaluated at 7 hpg (**Figure 1C, Figure S1**). In the FD10-gavaged controls, FD10 remained in the intestinal bulb and mid-intestine, migrating solely through the cloaca. In contrast, co-gavage with UFP perfused intestinal FD10 fluorescence into venous capillary plexus, between the dorsal aorta (DA) and posterior cardinal vein (PCV). Approximately 80 % of UFP-gavaged embryos exhibited luminal fluorescence in both AVP and CVP. In addition, co-gavage with EDTA that induced structural deformation of epithelial TJ phenocopied UFP-gavaged embryos (* *p < 0.05* vs. FD10, n= 10 per each group) (**Figure 1E-G**). We further examined whether acute UFP ingestion regulates villus ultrastructure in the embryonic GI tract. At 7 hpg, the distribution of intestinal FD10 fluorescence was assessed to confer the morphology of the mid-intestine (**Figure 2A**). While luminal FD10 fluorescence in mid-intestine was prominent in FD10-gavaged controls, co-gavage with UFP systemically reduced intestinal FD10 fluorescence in 80% of gavaged embryos (* *p < 0.05* vs. FD10, n= 10 per each group) (**Figure 2B-D**). Thus, our data suggest that acute UFP ingestion via micro-gavage disrupts the intestinal barrier integrity and retards maturation of the embryonic GI system.

**Figure 2.**
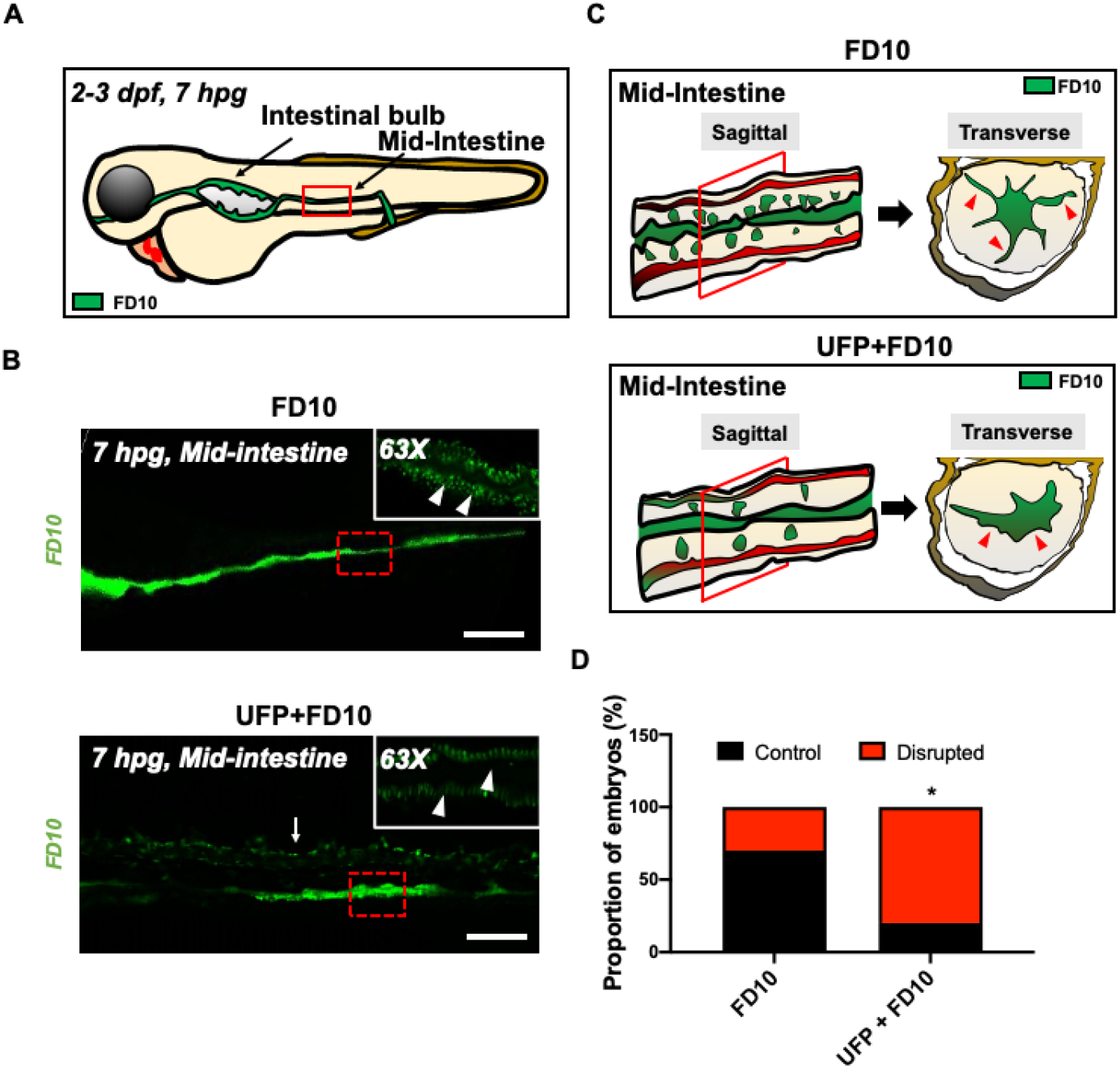
Acute UFP exposure disrupts maturation of embryonic GI tract. **(A)** Schematic representation of the embryonic GI tract at 7 hpg. The density of FD10 fluorescence in the mid-intestine was assessed to evaluate maturation of the GI tract (Red box). **(B)** Representative images of UFP-disrupted GI tract (white dashed box). Compared to FD10-gavaged controls, co-gavaging UFP, as denoted with endoluminal FD10 fluorescence (white arrow), altered morphology and systemically reduced the density of FD10 fluorescence in the mid-intestine (white arrowheads, n=5 per group). Scale bar: 20 μm. **(C)** Schematic representations of sagittal and transverse views of the mid-intestine with and without UFP gavage. Acute UFP exposure in developing GI system retards maturation (red arrowheads). **(D)** Percentage of embryos exhibiting reduced FD10 density in the mid intestine (\**p < 0.05* vs. FD10, n=10 per group).

### 3.2 Micro-angiography via CCV to mimic UFP gavage

The UFP-disrupted intestinal barrier is further supported by performing micro-angiography in the zebrafish micro-circulation system (**Figure 3A**). Immediately following micro-angiography, the micro-circulatory system in transgenic *Tg*(*flk1: mCherry*) embryos, including the injection site, CCV, heart, the DA and PCV, exhibited prominent FD10 fluorescence (**Figure 3B**). At 1 hour post injection (hpi), FD10 fluorescence was observed in both AVP and CVP mimicking UFP gavage (n=5) (**Figure 3C-D**). Thus, our micro-angiography results suggested the notion that UFP gavage impaired endothelial microenvironment via disrupted intestinal barrier.

**Figure 3.**
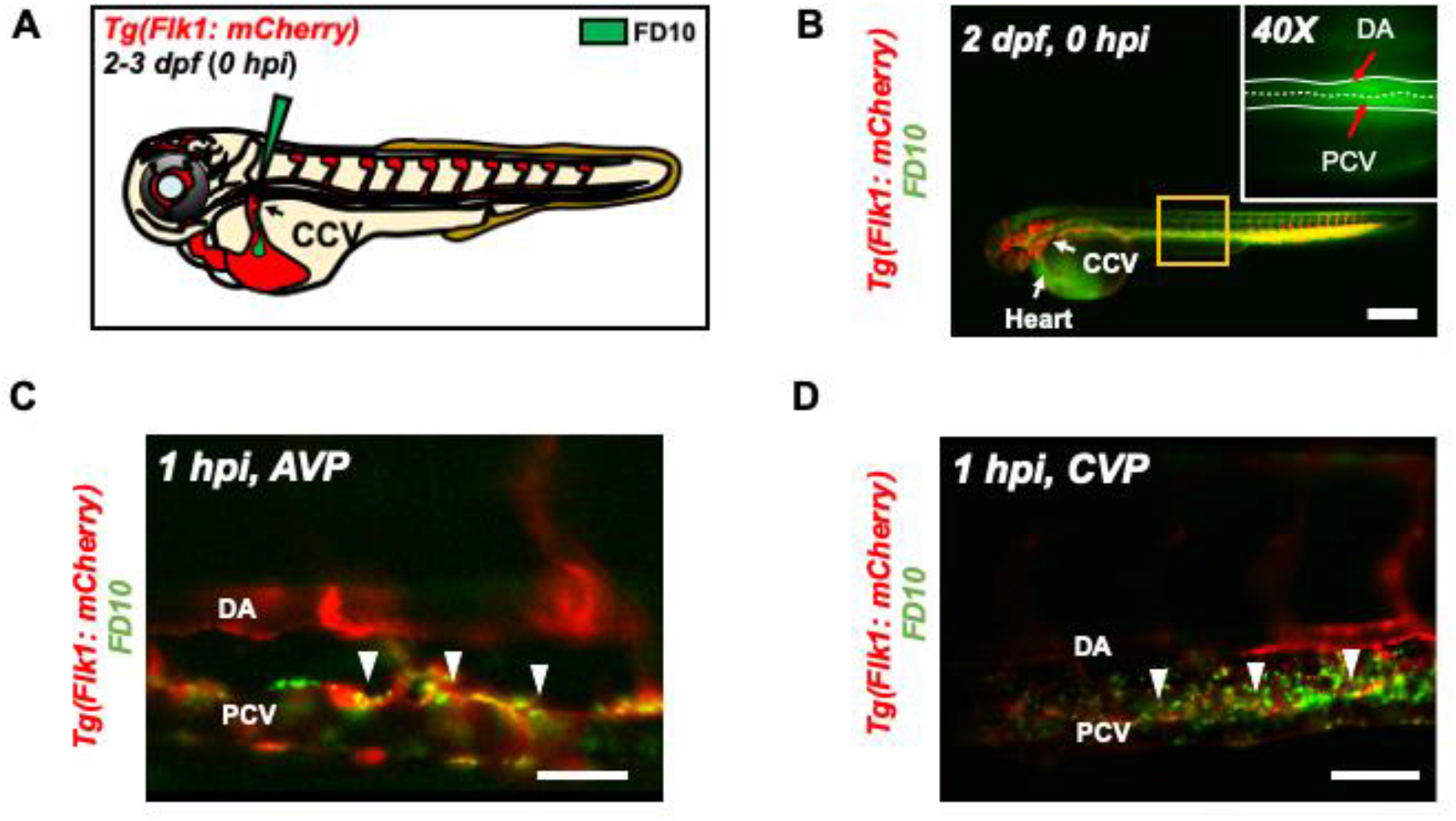
Micro-angiography via common cardinal vein (CCV) to mimic UFP gavage. **(A)** A schematic representation of micro-angiography via CCV to introduce FD10 to the microcirculatory system. **(B)** A representative image of the transgenic *Tg*(*flk1: mCherry*) embryo following FD10 injection to CCV. At 0 hpi, FD10 fluorescence was prominent at the injection site, CCV, heart, DA and PCV. Scale bar: 100 μm. **(C-D)** At 1 hpi, FD10 was distributed in the AVP and CVP, between the DA and PCV, mimicking UFP gavage-mediated effects (white arrowheads, n=5 per group). Scale bar: 32 μm.

### 3.3 UFP exposure regulates Notch-mediated endothelial TJ expression

To corroborate UFP-disrupted intestinal barrier, we assessed the transcription of endothelial TJ, including Zo1, Cldn1, and Ocln1 *in vitro*. Exposure to UFP decreased Zo1 and Cldn1 mRNA expression by 23% and 34%, respectively, whereas Ocln1 mRNA expression remained unchanged (*\* p < 0.05* vs. H2O, n=3). Treatment with the small molecular inhibitor Iwr1, to down-regulate Wnt-mediated TJ expression, reduced Zo1 mRNA by 37% and significantly inhibited Cldn1 and Ocln1 mRNA expression by 83% and 71%, respectively, whereas treatment with lithium chloride (LiCl) for ectopic activation of Wnt/ β-catenin signaling pathway and prevent loss of TJ upregulated both Cldn1 and Ocln1 mRNA expression by 3.1-fold and 2.8-fold (*\* p < 0.05* vs. DMSO for Iwr1, and vs. H2O for LiCl, n=3) (**Figure 4A**).[30,31] As a corollary, UFP exposure led to a dose-dependent reduction in Zo1 and Cldn1 mRNA expression. UFP at 50 μg·mL^-1^ (UFP50) attenuated Zo1 and Cldn1 mRNA expression by 22% and 12%, respectively, as compared to UFP at 25 μg·mL-1 (UFP25) (*\* p < 0.05* vs. H2O, n=3). Similarly, Hes1 mRNA, a Notch target gene, was also reduced in dose-dependent manner (**Figure 4B**). Treatment of Adam10 inhibitor to suppress global Notch receptor activation down regulated both Cldn1 and Hes1 mRNA expression by 22% and 69%, respectively, as compared to the untreated controls (*\* p < 0.05* vs. DMSO, n=3) (**Figure 4C**). This finding suggested Notch activity implicates UFP-mediated Cldn1 mRNA expression. Co-gavaging Adam10 inhibitor in zebrafish embryos mimicked UFP gavage by developing luminal FD10 fluorescence in both AVP and CVP. As a corollary, global overexpression of NICD mRNA via micro-injection restored UFP-disrupted intestinal barrier (*\* p < 0.05* vs. FD10, n=10 per group) (**Figure 4D, E**). Taken together, our data recapitulated UFP exposure modulates TJ expressions in Notch-dependent manner in association with impaired development of the GVB.

**Figure 4.**
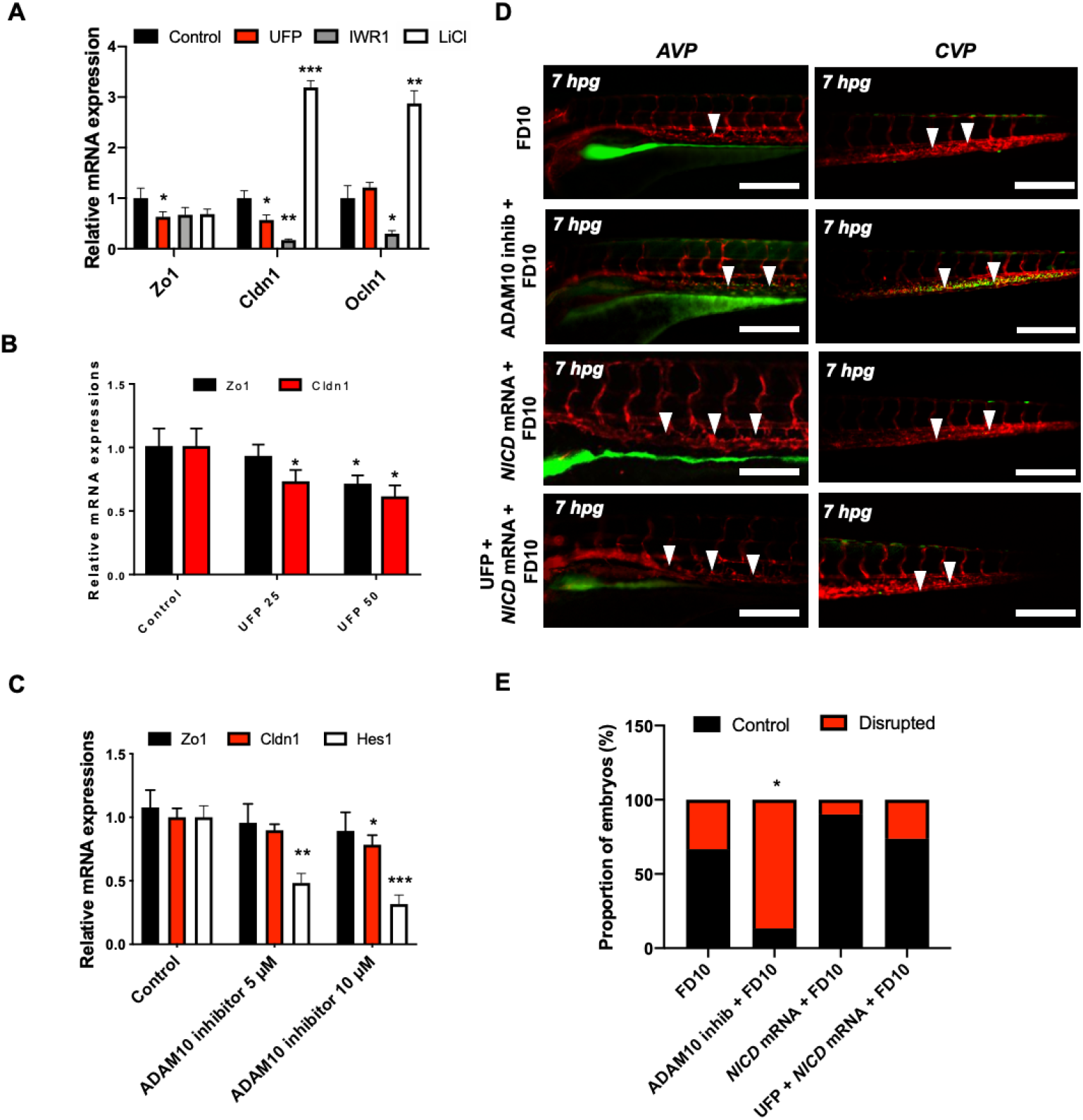
UFP exposure down-regulates mRNA expressions of endothelial tight junction (TJ) protein and the Notch target gene to disrupt the GVB. mRNA expressions of TJ proteins, including zonula occludens1 (Zo1), claudin 1 (Cldn1), and occludin 1 (Ocln1), and the Notch target genes, including Hairy and enhancer of split-1 (Hes1), were assessed *in vitro* by cultured human aortic endothelial cells (HAEC). **(A)** UFP exposure (25 μg·ml^-1^ for 6 hours) inhibited Zo1 and Cldn1 mRNA expressions, whereas Ocln1 mRNA remained unchanged. While Iwr1 treatment (10 μM) diminished overall TJ mRNA expression, LiCl (20 mM) up-regulated Cldn1 and Ocln1 mRNA expression (*\* p < 0.05* vs. DMSO for Iwr1, H2O for LiCl, n=3). **(B)** UFP exposure (25-50 μg·mL^-1^ for 6 hours) down-regulated both TJ (Zo1 and Cldn1 mRNA) and Notch target genes (Hes1) mRNA expression in a dose-dependent manner (* *p < 0.05* vs. H2O, n=3). **(C)** Treatment of Adam10 inhibitor (5 μM) to inhibit Notch receptor activation down-regulated Cldn1, and Hes1 mRNA in a dose-dependent manner (* *p < 0.05* vs. DMSO, n=3) **(D)** Micro-gavage with Adam10 inhibitor promoted transmigration of FD10 to both AVP and CVP. Micro-injection of NICD mRNA restored UFP- and Adam10 inhibitor-mediated effect. **(E)** Percentage of embryos exhibiting endoluminal FD10 fluorescence (* *p < 0.05* vs. FD10, n=10 per group).

### 3.4. Figures, tables and Schemes

**Figure S1.**
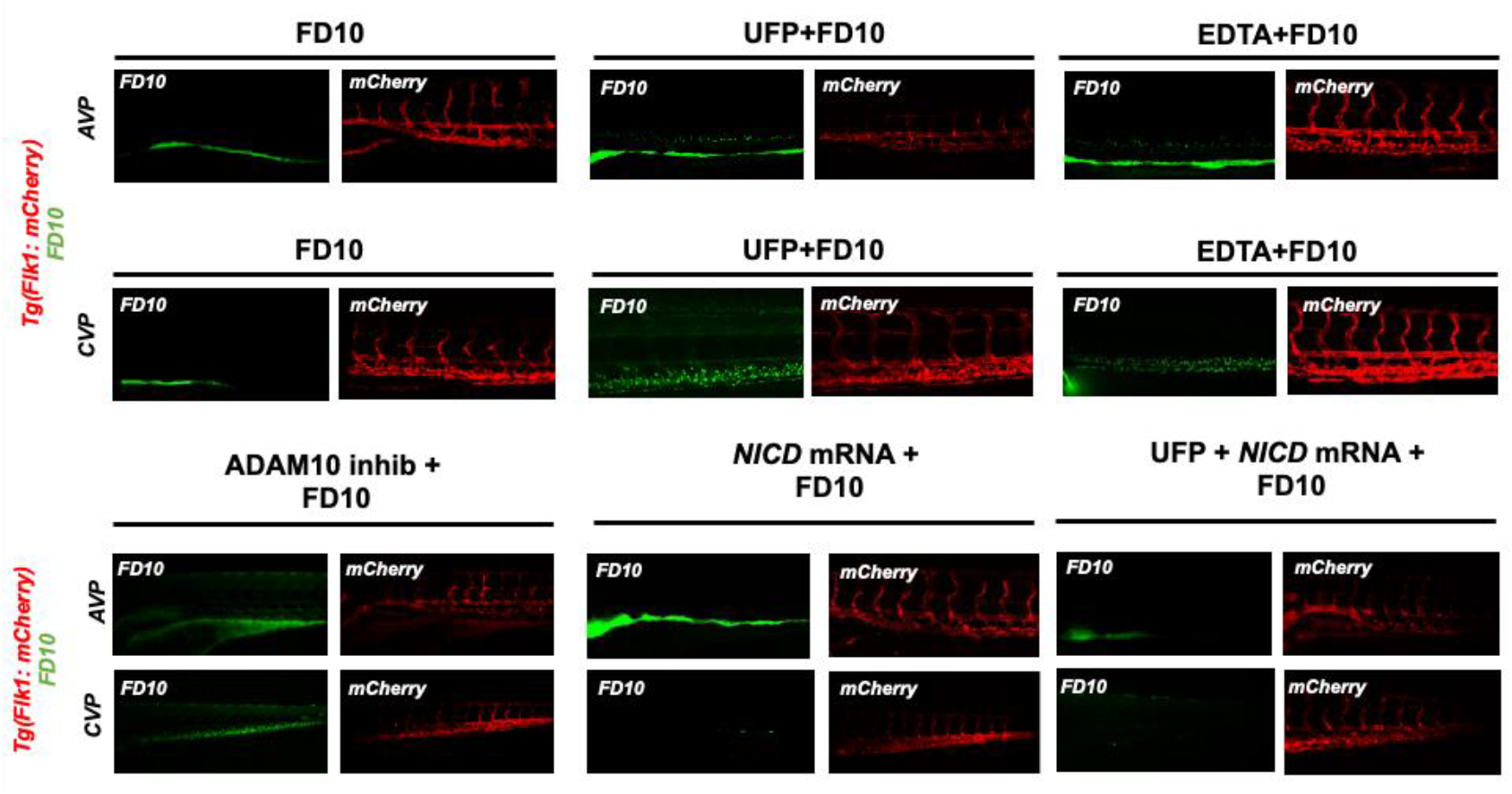
Imaging endoluminal distribution of FD10. Images of FD10 (green) and vascular endothelium (*flk1*^+^, red) were taken respectively in response to UFP, Adam10 inhibitor, and *NICD* mRNA rescue and superimposed to assess vascular endothelial distribution of FD10.

## 4. Discussion

The novel contribution of our study is to elucidate UFP-disrupted intestinal barrier integrity using the zebrafish system. Utilization of zebrafish embryos provided a high-throughput screening of altering gut-vascular permeability. Micro-gavage of ambient UFP to the transgenic zebrafish embryos promoted transmigration of FD10 from the intestinal epi-lumen to vascular endo-lumen (**Figure 1**). UFP exposure further led to an impaired embryonic villus ultrastructure during development (**Figure 2**). Micro-gavage of Adam 10 inhibitor disrupted GVB, whereas micro-injection of *NICD* mRNA rescued UFP-disrupted intestinal barrier. As a corollary, UFP exposure down-regulated Notch-mediated TJ expression in cultured endothelial cells (**Figure 4**). Overall, UFP exposure down-regulates Notch signaling-mediated TJ expression to increase endothelial permeability, and subsequently disrupted the GVB (**Figure 5**).

**Figure 5.**
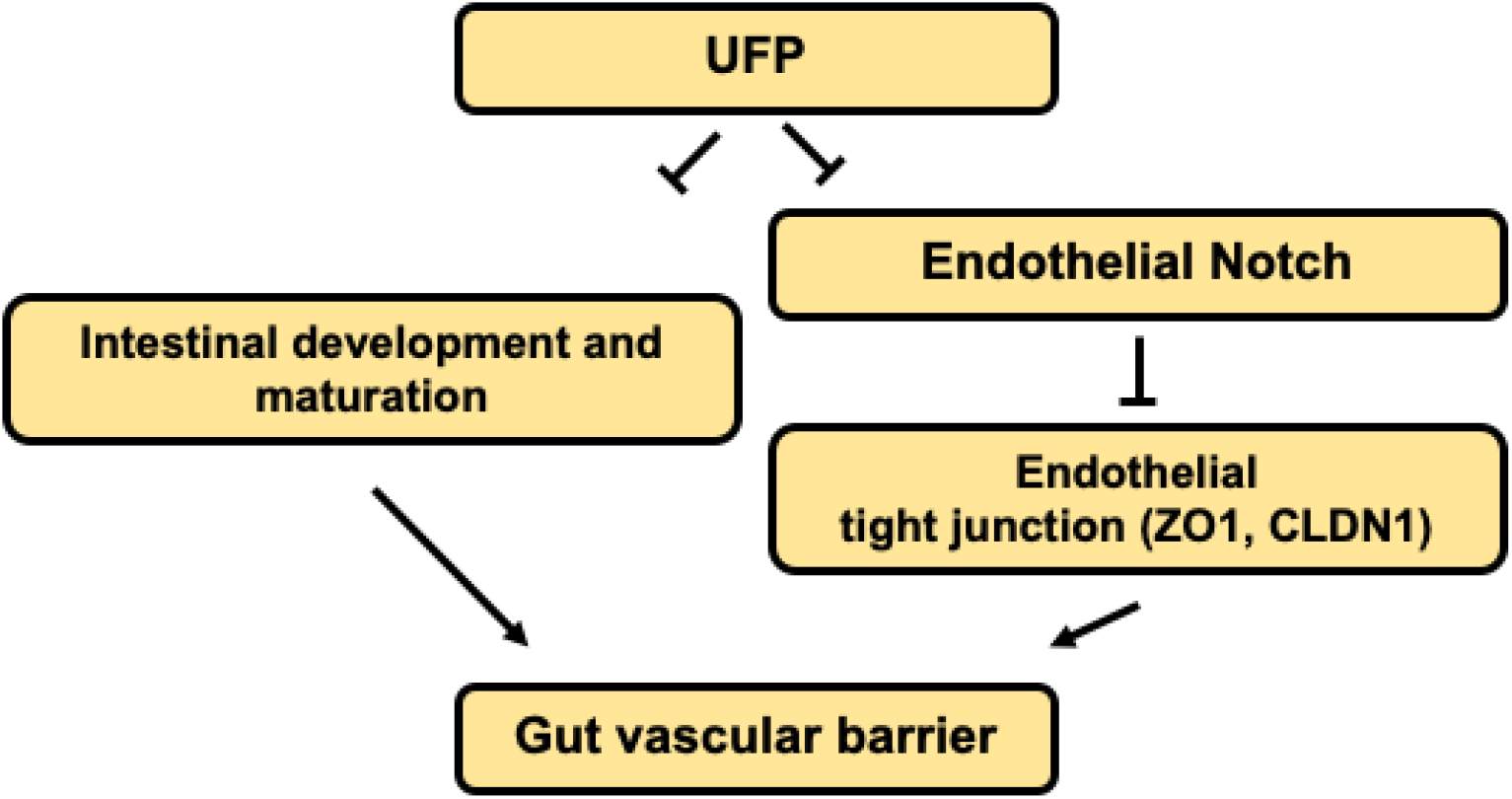
Schematic overviews of the proposed mechanisms

Ambient UFP are the redox-active sub-fraction of PM2.5, harboring elemental and polycyclic aromatic hydrocarbons emitted from primary diesel combustion and photochemical formation from urban environmental gases. [10] Their small size and large surface-to-volume ratio facilitates potential adsorption in the cardiopulmonary and vascular system associated with pathophysiology of systemic inflammatory responses.[32–34] Increasing epidemiological studies correlate UFP exposure with clinical relevance to intestinal disease and gut microenvironment. Inhaled or dietary UFP ingestion aggravates intestinal dysbiosis and macrophage infiltrates in the GI tract, suggesting altered intestinal barrier. [10,17,35] Herein, our integration of an embryonic zebrafish model with micro-gavage technique provides molecular insights into UFP exposure to disrupt the gut-vascular homeostasis.

Notch signaling is well-recognized as a conserved mechanism for GI epi- and endothelial homeostasis.[36,37] In developing gut, Notch signaling involves multi-potential stem/progenitor cell proliferation and lineage.[38] Inhibition of global Notch activity, including pharmacological inhibiton, genetic recombination, and neutralizing antibodies, leads to an overall reduction in gastric epithelial and intestinal stem cell proliferation, whereas overexpression of NICD to increase systemic Notch activity promotes the proliferation of gastric stem cells. [39–43] Moreover, intestinal Notch activation via Delta D programs the differentiation of absorptive enterocytes. [38] While reduction in Notch activity results in secretory cell hyperplasia, constitutively-active NICD shifts the differentiation of secretory cells.[38,44] The interplay between Wnt and Notch is also recognized to direct the fate of intestinal epithelial cells.[45] Dysregulation of the Notch ligand Delta-like ligand 4 (Dll4), or Notch 1 receptor expression, induces hyper-permeability and transcytosis in murine retina, while down-regulation of Notch 4 expression is associated with endothelial blood-brain barrier dysfunction.[46,47] The role of Notch activity in vascular stabilization and cell quiescence is also well-recognized.[37] In parallel, differential patterning of VE-cadherin in the absence of Notch activity supports the regulatory effects of junctional stability.[48] Consistent with the current literature, our data supports that acute UFP ingestion inhibits Notch-mediated intestinal barrier integrity during development (**see the proposed mechanism in Figure 5**).

The transcription factor Forkhead box sub-family O1 (FOXO1) enhances repressor clearance and forms a transcriptional activation complex during Notch activation.[24,49] In colonic tumorous tissue, ectopic level of FOXO1 expression alters epithelial permeability, villus ultrastructure, and arrangements in the GI tract.[50] Furthermore, intestinal epithelial FOXO1 expression associates GI barrier integrity.[51,52] At the molecular level, concurrent nuclear retention of FOXO1 and β-catenin represses Cldn5 expression in the cerebral vascular endothelium.[53] While ambient PM exposure epigenetically controls of cardiometabolic state, endothelial FOXO1 couples cellular metabolism and endothelial homeostasis.[54] Our previous report indicates UFP exposure-mediated cytoplasmic FOXO1 expressions in vascular endothelial cells.[24] Whether acute UFP ingestion decreases intestinal FOXO1 expression to participate in Notch-mediated GVB warrants further investigation.

3-dimensional (3D) imaging techniques with high spatiotemporal resolution to capture the dynamics of GI tract remain as an imaging challenge. The advent of light-sheet fluorescence microscopy (LSFM) allows for real-time imaging of the peristaltic contraction and the ultrastructure of the GI tract. [55–57] The integration of LSFM with zebrafish genetics and deep learning for post-imaging processing enables precise time-lapse monitoring of myocardial contraction and intracardiac flow dynamics. [58] Thus, interfacing LSFM with the transgenic zebrafish system and micro-gavage technique may allow for simultaneous 3D imaging of the FD10 distribution and disrupted intestinal barrier to unravel epigenetic cues underlying UFP exposure in the developing GI tract.

Overall, we demonstrate a zebrafish model for rapid screening of GVB in response to epigenetic stimuli. We demonstrate new molecular insights into UFP-mediated Notch signaling to disrupt GVB.

## Supplementary Materials

Figure S1. Imaging endoluminal distribution of FD10

## Author Contributions

Conceptualization, Baek.KI. and Hsiai.TK.; methodology, Baek.KI. and Qian.YI.; software, Baek.KI. and Chang.CC.; validation, Chang.CC. and O’Donnell.R.; formal analysis, Baek.KI.; investigation, Baek.KI.; resources, Soleimanian.E., Sioutas.C. and Hsiai.TK.; writing-original draft preparation, Baek. KI. And Qian.YI.; writing-review and editing, Hsiai. TK. and O’Donnell. R.; visualization, Baek.KI. and Chang. CC.; supervision, Hsiai.TK.; project administration, Hsiai.TK.; funding acquisition, Hsiai, TK.

“All authors have read and agreed to the published version of the manuscript.”

## Acknowledgments

The authors like to express their gratitude to Dr. Weinmaster for providing the NICD plasmid, and Mrs. Yuan Dong at UCLA zebrafish core facility for generously providing *Tg*(*flk1:mCherry*) zebrafish line.

